# Mechanically activated bone cells drive vessel formation via an extracellular vesicle mediated mechanism

**DOI:** 10.1101/2023.02.10.527969

**Authors:** N. Shen, M. Maggio, I. Woods, M. Lowry, K.F Eichholz, E. Stavenschi, K. Hokamp, F.M. Roche, L. O’Driscoll, D.A. Hoey

## Abstract

Blood vessel formation is an important initial step for bone formation during development as well as during remodelling and repair in the adult skeleton. This results in a heavily vascularized tissue where endothelial cells and skeletal cells are constantly in crosstalk to facilitate homeostasis, a process that is mediated by numerous environment signals, including mechanical loading. Breakdown in this communication can lead to disease and/or poor fracture repair. Therefore, this study aimed to determine the role of mature bone cells in regulating angiogenesis, how this is influenced by a dynamic mechanical environment, and understand the mechanism by which this could occur. Herein, we demonstrate that both osteoblasts and osteocytes coordinate endothelial cell proliferation, migration, and blood vessel formation via a mechanically dependent paracrine mechanism. Moreover, we identified that this process is mediated via the secretion of extracellular vesicles (EVs), as isolated EVs from mechanically stimulated bone cells elicited the same response as seen with the full secretome, while the EV depleted secretome did not elicit any effect. Despite mechanically activated bone cell derived EVs (MA-EVs) driving a similar response to VEGF treatment, MA-EVs contain minimal quantities of this angiogenic factor. Lastly, a miRNA screen identified mechanoresponsive miRNAs packaged within MA-EVs which are linked with angiogenesis. Taken together, this study has highlighted an important mechanism in osteogenic-angiogenic coupling in bone and has identified the mechanically activated bone cell derived EVs as a therapeutic to promote angiogenesis and potentially bone repair.

## Introduction

It is well established that the vasculature is crucial for bone development and remodelling as it is the main source of oxygen, hormones and growth factors delivered to resident cells within skeletal tissue [1]. The vasculature in bone is formed predominately via angiogenesis, the process by which new vessels sprout and grow from pre-existing vessels [2]. The process of angiogenesis and the resulting vessels are specialized to bone (Type H and L capillaries) and are regulated by numerous microenvironmental signals [3]. For instance, mature bone cells secrete pro-angiogenic factors such as vascular endothelial growth factor (VEGF) that can induce angiogenic responses in resident endothelial cells. Endothelial cells in turn also release factors that can regulate chondrogenesis, osteogenesis, and maintain the hematopoietic stem cell niche within the bone marrow [4]. In addition to maintaining homeostasis, angiogenesis plays an important role in fracture repair and regeneration [5]. It is therefore, not surprising that defects in the formation or regeneration of the skeletal vasculature can be a major contributor to numerous pathologies, including tissue necrosis, osteoporosis, and cancer [6]. Understanding the intricate relationship between angiogenesis and osteogenesis may reveal new insights into bone (patho)physiology and open new avenues to treat skeletal disease and regenerate defects.

A potent regulator of new bone formation in development, remodelling, and during fracture repair is mechanical loading [7-9]. This loading has similarly been shown to regulate angiogenesis [10], although it is unclear whether these are independent or related effects. Cells of the osteogenic lineage can release factors that regulate endothelial cell proliferation, migration, and angiogenesis [2]. The pro-angiogenic effect of the mesenchymal stem/stromal cell (MSC) secretome is well established and is a driving force for many cell therapies utilizing this cell type for tissue repair [11]. Interestingly, the pro-angiogenic properties of the MSC secretome is enhanced following mechanical loading [12]. The angiogenic properties of this secretome is also modified as the MSC undergoes osteogenic lineage commitment [13]. Both the osteoblast and osteocyte have also been shown to regulate angiogenesis [14, 15], indicating that cells of the osteogenic lineage may coordinate blood vessel formation. Moreover, while mechanical activation of the osteoblast further enhances the angiogenic properties of the secretome [2], it is unclear if osteocytes also possess the same mechanically driven responses. Given the abundance of osteocytes in bone and the established role of the osteocyte network as essential transducers of mechanical signals [16-19], it is very likely that the osteocyte angiogenic properties are also mechanically regulated, although this has not been demonstrated to date.

Extracellular vesicles (EVs) are a group of highly heterogeneous cell-derived lipid based structures, which are involved in multiple physiological and pathological processes [20, 21]. They can interact with local and distant cellular targets and mediate their phenotype by transferring contents, which include varieties of functional lipids, proteins, and nucleic acids such as miRNAs, from one cell to another [21]. As such EVs have emerged as potent mediators of cell-to-cell communication in many tissues including bone tissue [22, 23]. For instance, MSC and osteoblast-derived EVs can act as a delivery vehicle for pro-angiogenic and pro-osteogenic paracrine factors [22, 24-26] and are being utilized as a cell-free therapy to enhance regeneration [27, 28]. More recently, osteocytes have been shown to release pro-osteogenic EVs and the regenerative potency of these EVs was dependent on the mechanical activation of the parent cell [29-31]. However, the angiogenic properties of both the osteoblast and osteocyte derived-EVs remain unknown. EVs therefore represent a potential mechanism by which mature bone cells may mediate angiogenesis.

Therefore, the aim of this study was to first, determine whether mature bone cells (osteoblasts and osteocytes) can regulate endothelial cell proliferation, migration, and angiogenesis and whether this process is mechanically regulated. Secondly, we aim to investigate whether this potential pro-angiogenic paracrine signal could be mediated via the secretion of extracellular vesicles, and lastly, we aim to explore the potential mechanism by which EVs may mediate an angiogenic response via the packaging and delivery of pro-angiogenic cargo. The identification of such would represent further insight into the complex coupling of osteogenesis and angiogenesis in bone and indicate the potential of bone derived EVs as a novel angiogenic therapeutic.

## Materials and methods

### Cell culture

Two cell lines and one primary cell type were used in this study; the MLO-Y4 osteocyte cell line (Kerfast); the MC3T3-E1 osteoblast cell line (ATCC) and human umbilical vein endothelial cells (HUVECs, Lonza). MLO-Y4 cells were maintained in α-MEM supplemented with 5% fetal bovine serum (FBS), 5% calf serum (CS) and 1% penicillin-streptomycin (P/S). MC3T3-E1 cells were maintained in α-MEM supplemented with 10% FBS and 1% P/S. HUVECs were maintained in EBM-2 MV BulletKit medium (CC-3156 and CC-4147; Lonza). HUVECs in passages 3-5 were used for all experiments.

### Mature bone cell conditioned medium

Osteocytes were seeded into collagen-coated 6-well-plates with a density of 60,000 cells per well and osteoblasts were seeded into 6-well-plates with a density of 80,000 cells per well. After 24 hours the cells were washed twice with PBS and subsequently 1.83 mL of serum-free α-MEM medium was added to each well. To mechanically stimulate cells, a fluid flow-induced shear stimulus was provided by an orbital shaker. Orbital shakers have previously been utilised as a simple and effective means of producing shear stimulus in circular plates [32] and have advantages over traditional parallel plate flow chambers in terms of the volume of media collection post-stimulus. This study was carried out using an orbital shaker model of shear stress application previously developed by Salek *et al* [33], who found that a frequency of 100 rpm applied to a 6-well plate could be used to produce a shear distribution of approximately 0-3.2 dyne/cm^2^ (0-0.32 Pa) (Fig. 1) [33]. Cells cultured statically in 6-well plates were used as static controls. After 24 hours, conditioned media (CM) was collected from static or mechanically stimulated osteoblasts or osteocytes and analyzed immediately or stored at -80 ºC for further analysis.

**Figure 1:**
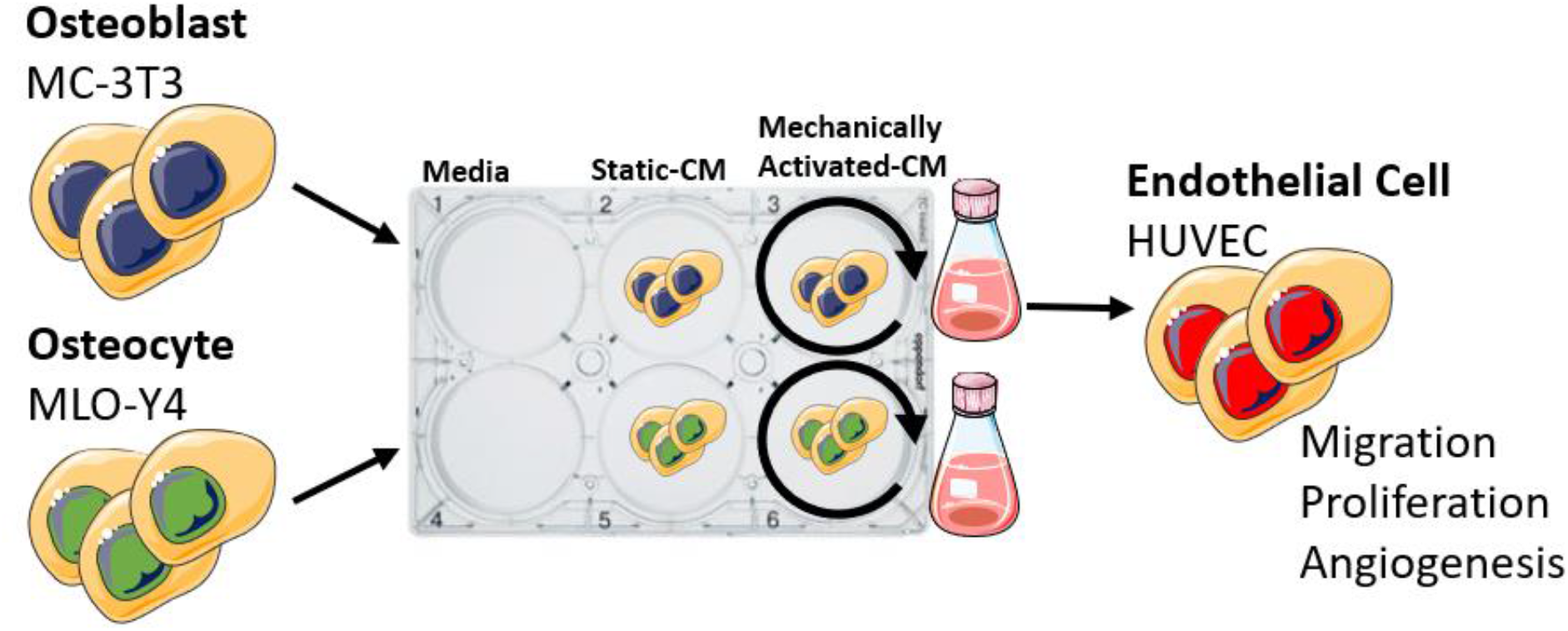
Schematic illustrating experimental design. Conditioned media (CM) was collected from osteoblasts and osteocytes cultured statically or subjected to dynamic fluid shear using an orbital shaker. CM was used to treat human endothelial cells and migration, proliferation, and tube formation was assayed.

### Extracellular vesicle collection

Extracellular vesicle (EV) collection was performed with an ultracentrifugation method as previously described [34]. In brief, CM was centrifuged at 300 g for 10 min and 2000 g for 15 min at 4 ºC to eliminate cells and debris, then filtered through a 0.45 μm pore filter. Subsequently, the medium was ultracentrifuged at 110,000 g for 75 min at 4 ºC. EV pellets were washed in PBS and centrifuged at 110,000 g for 75 min at 4 ºC again. Then EV pellets were resuspended in PBS and characterised by nanoparticle tracking analysis (NanoSight) and transmission electron microscope (TEM).

### Proliferation Assay

HUVECs were seeded onto 48-well plates at 40,000 cells per well. After 24 hours of incubation, 0.5 ml of control media (serum-free α-MEM medium, with and without 10 ng/mL VEGF supplement), mature bone cell conditioned media, or EV reconstituted medium was added in a 1:1 ratio with the serum-free EBM-2 medium. VEGF, a proangiogenic factor, was used as a positive control in all assays. After 24 hours of incubation, the wells were stained with DAPI. Fluorescent images were taken from each sample and the cell nuclei in each field were counted using ImageJ (NIH, Bethesda, MD).

### Migration assay

HUVECs were seeded onto 24-well cell culture inserts containing membranes with 8 μm pores (Millipore) at 10,000 cells per insert. After 4 hours of incubation, control media (serum-free α-MEM medium, with and without 10 ng/mL VEGF supplement), mature bone cell conditioned media, or EV reconstituted medium was mixed in 1:1 ratio with serum-free EBM-2 medium and added to the well. An equal volume of serum-free EBM-2 medium was added to the top of the inserts. The cells were incubated for 18 hours, and then the cells on the topside of the membranes were removed with a cotton swab. The remaining cells on the underside of the membranes were stained with hematoxylin. Nine brightfield images were taken from each sample with a brightfield microscope (Olympus).

### Tube formation assay

The tube formation assay was performed as previously described [2]. 48-well-plates were coated with Matrigel (growth factor reduced, Corning) at 100 μL per well and allowed to polymerize for 45 min at 37 ºC. HUVECs were seeded onto Matrigel-coated 48-well-plates at 40,000 cells per well. Control media (serum-free α-MEM medium, with and without 10 ng/mL VEGF supplement), mature bone cell conditioned media, or EV reconstituted medium was added in 1:1 ratio with serum-free EBM-2 medium to the wells. HUVECs were incubated for 18 hours to allow for tubule formation. Subsequently, the samples were fixed and stained with phalloidin and DAPI. For each sample, 3 fields of view were taken with a fluorescent microscope. The total tube length, the junction density, and the number of branches were quantified using ImageJ (National Institute of Health, USA) and normalized to the negative control.

### Nanoparticle tracking analysis

Nanoparticle tracking analysis was performed on EVs with the NTA NS500 system (NanoSight, Amesbury, UK) to determine particle size based on Brownian motion. EV samples were diluted at 1:50 in PBS and injected into the NTA system, which obtained four 40-second videos of the particles in motion. Videos were then analyzed with the NTA software to determine particle size.

### Transmission electron microscopy

A 20 μL aliquot of EVs was placed onto parafilm (Sigma–Aldrich). A 300-mesh copper grid (Agar Scientific) was placed on top of the drop for 45 min. The grid was subsequently washed three times in 0.05 M phosphate buffer (freshly prepared using dihydrogen potassium phosphate (Sigma–Aldrich) and dipotassium hydrogen phosphate (Merck)) for 5 min, fixed in 3% glutaraldehyde (Agar Scientific) for 10 min, washed three times for 5 min in dH2O and contrasted in 2% uranyl acetate (BDH). Grids were examined at 100 kV using a JEOL JEM-2100 TEM.

### ELISA

ELISA kits for mouse VEGF (R&D Systems, UK) were used according to the manufacturer’s instructions. 100 μL of mature bone cell CM or EV reconstituted medium were tested per well. Values were assayed in triplicate and calibrated against a VEGF standard. The sensitivity of the assay is 15.6 pg/mL and it detects both VEGF_120_ and VEGF_164_.

### miRNA library construction and high throughput sequencing

Total RNA was extracted using Trizol reagent (Invitrogen, CA, USA) following the manufacturer’s procedure. The total RNA quality and quantity was analyzed with Bioanalyzer 2100 (Agilent, CA, USA) with RIN number >7.0. Approximately 1 ug of total RNA was used to prepare small RNA library according to protocol of TruSeq Small RNA Sample Prep Kits (Illumina, San Diego, USA). Single-end sequencing 50bp was performed on an Illumina Hiseq 2500 at the LC Sciences (Hangzhou, China) following the vendor’s recommended protocol.

### Bioinformatics analysis of miRNA-seq data

Raw reads were subjected to a proprietary program, ACGT101-miR (LC Sciences, Houston, Texas, USA) to remove adapter sequences, low quality reads, common RNA families (rRNA, tRNA, snRNA, snoRNA) and repeats. Subsequently, unique sequences with length in 18–26 nucleotides were mapped to mouse precursors in miRBase 22.0 by BLAST search to identify known mouse miRNAs [35]. Length variations at both 3′ and 5′ ends and one mismatch inside the sequence were accepted in the alignment. Data quality was assessed using FastQC [36]. miRNAs in static and mechanically stimulated samples were profiled in three biological replicates. Differential expression analysis between conditions was conducted using DESeq2 [37]. Only miRNAs with read counts >25 in two or more replicates in at least one of the treatment groups were included in the analysis. The threshold for significance was set to p-adjusted value <= 0.05 (Benjamini-Hochberg method) and fold change >= 2. Gene ontology enrichment analysis of the miRNAs detected in the exosomes was carried out using the miRNA enrichment analysis and annotation tool, miEAA, (https://ccb-compute2.cs.uni-saarland.de/mieaa2/) [38]. miRNA gene targets were predicted from TargetScan (http://www.targetscan.org/) and subsequent gene ontology and KEGG pathway enrichment analysis of these targets was carried out using clusterProfiler (version 4.6) [39].

### Data analysis

Proliferation assay, migration assay and tube formation assay were normalized to no VEGF sample and were analysed using a one-way ANOVA, with Bonferroni post-hoc tests. All other analyses were performed using two-tailed unpaired student’s t-test with Wilcoxon correction. All data were analysed using GraphPad Prism 5. Statistically significant differences were indicated as *p<0.05.

## Results

### Mechanically stimulated osteoblasts and osteocytes secrete paracrine signals that promote human endothelial cell proliferation and recruitment

To determine if mature bone cells can coordinate angiogenesis, we first examined HUVEC proliferation and recruitment in response to conditioned media collected from statically cultured and mechanically activated osteoblasts and osteocytes (Fig.2).

**Figure 2:**
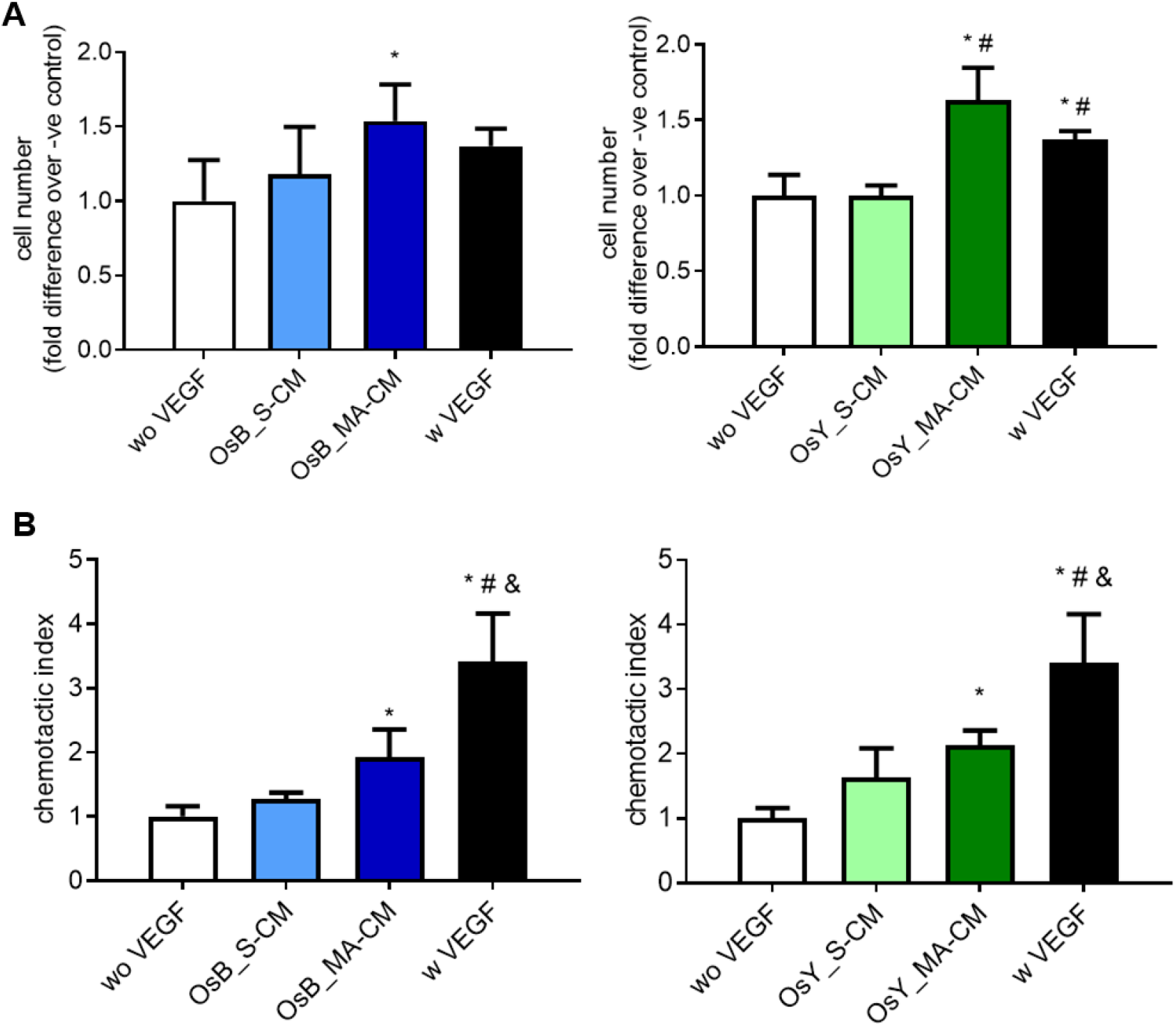
Mechanically activated osteoblasts and osteocytes promote HUVEC proliferation and migration via a paracrine mechanism. **(A)** Quantification of HUVEC number after 24 hours cultured in fresh medium without VEGF (negative control), with VEGF (positive control), statically (OsB_S-CM) or mechanically activated (OsB_MA-CM) MC3T3-E1 CM, and statically (OsY_S-CM) or mechanically activated (OsY_MA-CM) cultured MLOY4 CM. **(B)** Quantification of HUVEC migrated through a porous membrane in fresh medium without VEGF (negative control), with VEGF (positive control), statically or mechanically activated MC3T3-E1 CM, and statically or mechanically activated MLOY4 CM. Data presented as Mean ± SD, N = 3-6. * p<0.05 VS wo VEGF, # p<0.05 VS S-CM, & p<0.05 VS MA-CM.

The proliferation of endothelial cells was not influenced by conditioned media collected from statically cultured osteoblasts or osteocytes (S-CM) (Fig.2A). However, following mechanical stimulation of the mature bone cells, endothelial cell proliferation was significantly increased 1.5-fold (p<0.05) and 1.6-fold (p<0.05) when treated with mechanically activated conditioned media (MA-CM) from osteoblasts and osteocytes respectively, when compared to no VEGF media controls. Moreover, this increase in proliferation was similar to treatment with media containing 10ng/ml VEGF. The recruitment of endothelial cells was similarly influenced by osteoblasts and osteocytes. Conditioned media from statically cultured osteoblasts and osteocytes elicited a small non-significant 1.3-fold and 1.6-fold increase in endothelial cell migration (Fig.2B). However, as seen with proliferation, following mechanical activation of osteoblasts and osteocytes, a significant 1.9-fold (p<0.05) and 2.1-fold (p<0.05) increase in endothelial recruitment was identified respectively when compared to no VEGF media controls (Fig.2B). While MA-CM elicited a similar effect on endothelial proliferation as VEGF, in terms of recruitment, VEGF significantly outperformed CM from both types of mature bone cells.

Taken together, this data demonstrates that paracrine factors released from mature bone cells subjected to mechanical stimulation can enhance the proliferation and recruitment of endothelial cells in preparation for early angiogenesis.

### Mechanically stimulated osteoblasts and osteocytes secrete paracrine signals that coordinate angiogenesis

To determine whether mature bone cells can coordinate angiogenesis, we next examined HUVEC tubule formation in response to conditioned media collected from statically cultured and mechanically activated osteoblasts and osteocytes (Fig.3).

**Figure 3:**
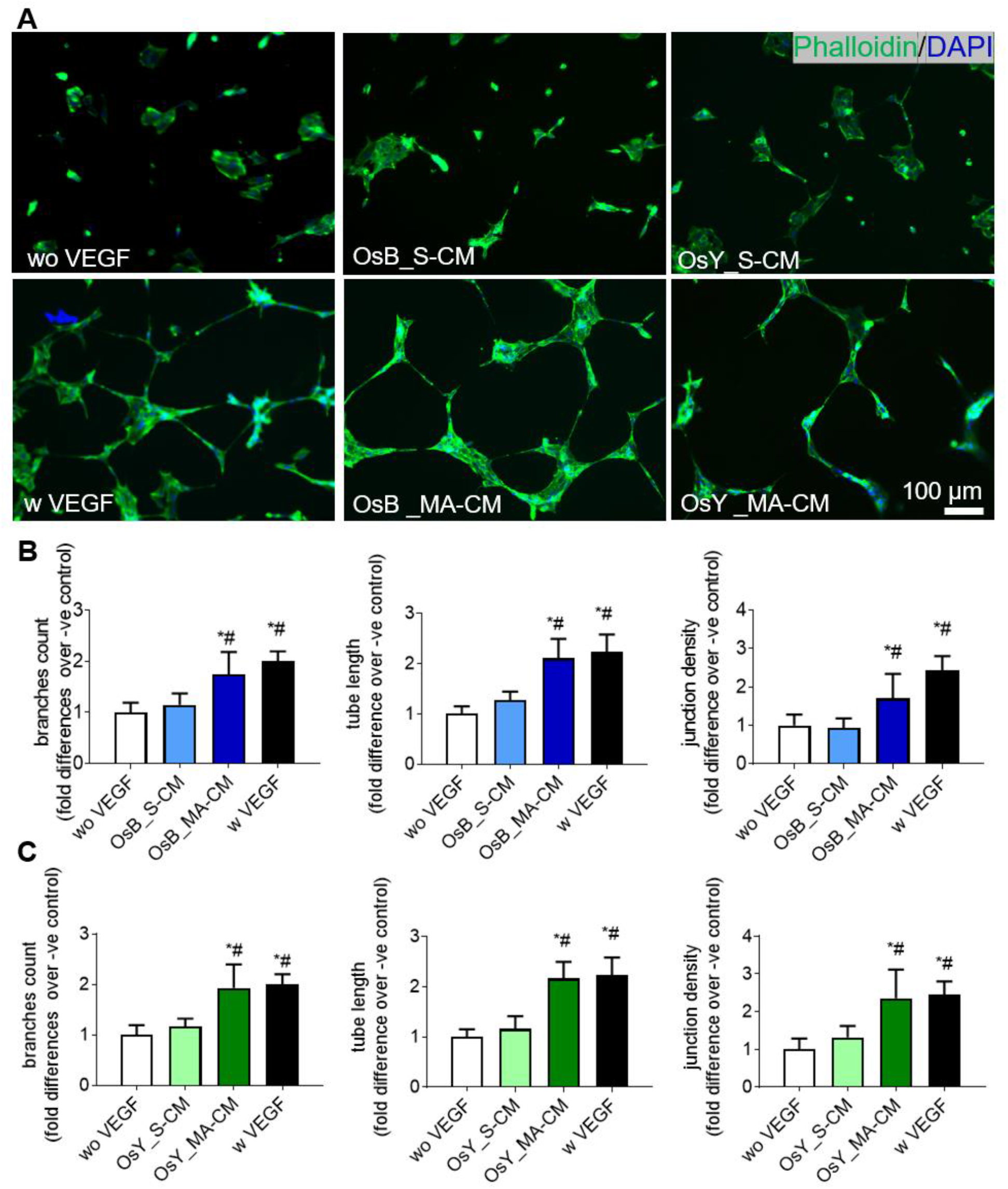
Mechanically activated osteoblasts and osteocytes induce vessel formation in HUVECs via a paracrine mechanism. **(A)** HUVECs cultured on Matrigel in fresh medium without VEGF (negative control), with 10 ng/mL VEGF (positive control), treated with conditioned medium derived from statically cultured MC3T3-E1 (OsB_S-CM), mechanically activated MC3T3-E1(OsB_MA-CM), statically cultured MLOY4 (OsY_S-CM), and mechanically activated MLOY4 (OsY_MA-CM). **(B)** Quantification of number of branches, tube length, junction density of tubes formed by HUVECs treated with conditioned medium derived from statically or mechanically activated MC3T3-E1. **(C)** Quantification of number of branches, tube length, junction density of tubes formed by HUVECs treated conditioned medium derived from statically or mechanically activated MLO-Y4s. Data presented as Mean ± SD, N = 3-7. * p<0.05 VS wo VEGF, # p<0.05 VS S-CM.

A standard HUVEC tube formation assay was performed on Matrigel and following 18hrs treatment with 10ng/ml of VEGF clear tubule-like formation was evident when compared to no VEGF controls (Fig.3A). Upon quantification of these images, VEGF treatment was found to significantly enhance the formation of branches (2-fold, p<0.05), tube length (2.2-fold, p<0.05), and junction densities (2.4-fold, p<0.05), demonstrating the ability of this cell type to undergo angiogenesis as previously described (Fig.3B,C) [2].

We next supplemented HUVEC media 1:1 with CM collected from statically cultured osteoblasts and osteocytes and analysed tube formation. No evidence of angiogenesis was evident after 18hrs in both statically cultured osteoblast and osteocyte CM supplemented groups (Fig.3A) and this was confirmed following quantification of branches, tube length, and junction density (Fig.3B,C). Interestingly, following supplementation with mechanically activated osteoblast and osteocyte CM, tube formation was evident in both groups and mirrored that seen with VEGF supplementation (Fig.3A). Furthermore, quantification of these images revealed a statistically significant increase in the number of branches, tube length, and junction density resulting from treatment with mechanical activated mature bone cell CM when compared to both no VEGF negative controls and statically cultured bone cell CM (Fig.3 B, C). These data demonstrate that mature bone cells drive angiogenesis via a paracrine mechanism following mechanical stimulation.

### Mechanically stimulated osteoblasts and osteocytes secrete extracellular vesicles that promote human endothelial cell proliferation and recruitment

As extracellular vesicles are known to be potent mediators of cell-to-cell communication, we next investigated whether EVs may play a role in the mature bone cell regulation of angiogenesis following mechanical stimulation (Fig.4A). EVs were collected and characterized from CM from mechanically activated osteoblasts and osteocytes using an ultracentrifugation method. EVs displayed the typical cup-shaped morphology as shown in TEM images (Fig. 4B). The TEM images further confirmed the expected EV size. The range of EVs collected from osteoblast CM was between 50 to 300 nm and EVs collected from osteocytes CM were distributed in the range of 80-350 nm (Fig. 4D), indicating that both EVs collected from osteoblast CM and osteocyte CM contain vesicles consistent with exosomes and microvesicles.

**Figure 4:**
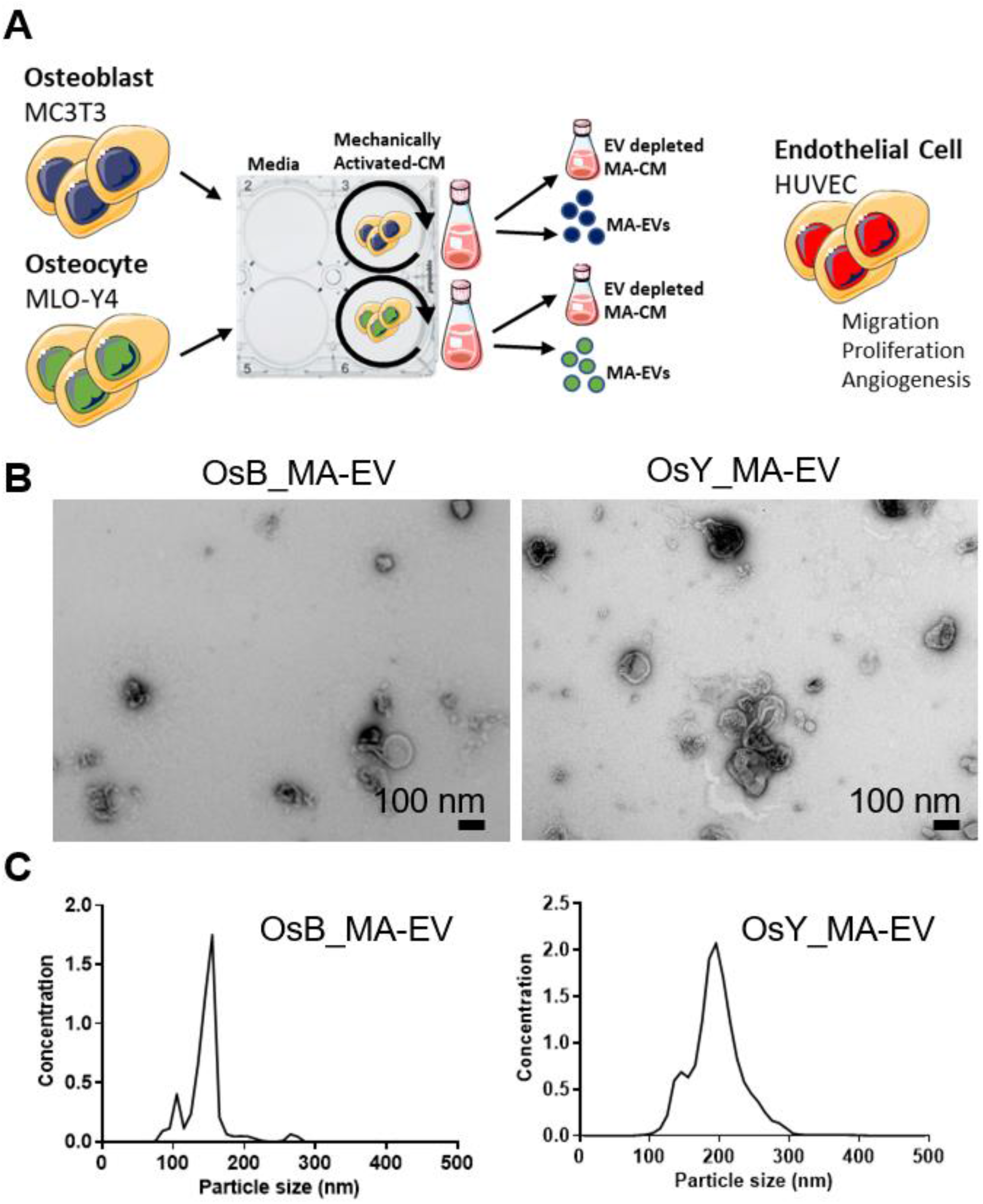
Characterisation of mechanically activated bone cell-derived extracellular vesicles (MA-EVs). **(A)** Conditioned media (CM) was collected from osteoblasts and osteocytes subjected to dynamic fluid shear using an orbital shaker. EVs were collected from mechanically activated CM (MA-CM). EV depleted MA-CM, along with isolated MA-EVs, were used to treat human endothelial cells and migration, proliferation, and tube formation was assayed. **(B)** TEM image of EVs isolated from mechanically activated osteoblasts (OsB_MA-EV) and osteocytes (OsY_MA-EV) **(C)** Nanoparticle size analysis on isolated EVs.

Our previous data demonstrated that only mechanically activated osteoblasts and osteocytes secreted paracrine factors that promoted endothelial cell proliferation, migration and angiogenesis. As such, the following study only focused on mechanically activated bone cell CM and the extracellular vesicles contained within. To assess the potential effect of mechanically activated bone cell derived EVs (MA-EVs) on angiogenesis, HUVECs were treated with EVs isolated from mechanically activated osteoblast and osteocyte CM and resuspended in the same volume of media. Furthermore, HUVECs were also treated with MA-CM which was depleted of EVs to ascertain whether soluble factors within the media may elicit the angiogenic response (Fig.4A)

Interestingly, osteoblast and osteocyte MA-CM, which previously enhanced endothelial cell proliferation, did not elicit any significant response when this media was depleted of extracellular vesicles (Fig. 5A). However, utilizing the EVs isolated from mechanically activated osteoblast and osteocyte-CM, a significant 2.6-fold (p<0.05) and 2.1-fold (p<0.05) increase in endothelial proliferation is seen (Fig.5A), which is consistent with that following 10ng/ml VEGF and mirrors that seen with MA-CM seen previously (Fig.2A). In terms of the recruitment of endothelial cells, MA-CM depleted of EVs from both osteoblasts and osteocytes elicited a small non-significant 1.3-fold and 1.6-fold increase in endothelial cell migration respectively, when compared to no VEGF controls (Fig.5B). EVs isolated from osteoblast MA-CM elicited a more robust 2.3-fold increase (p=0.052). However, EVs isolated from osteocyte MA-CM did not further influence endothelial cell recruitment when compared to EV-depleted MA-CM or VEGF controls.

**Figure 5:**
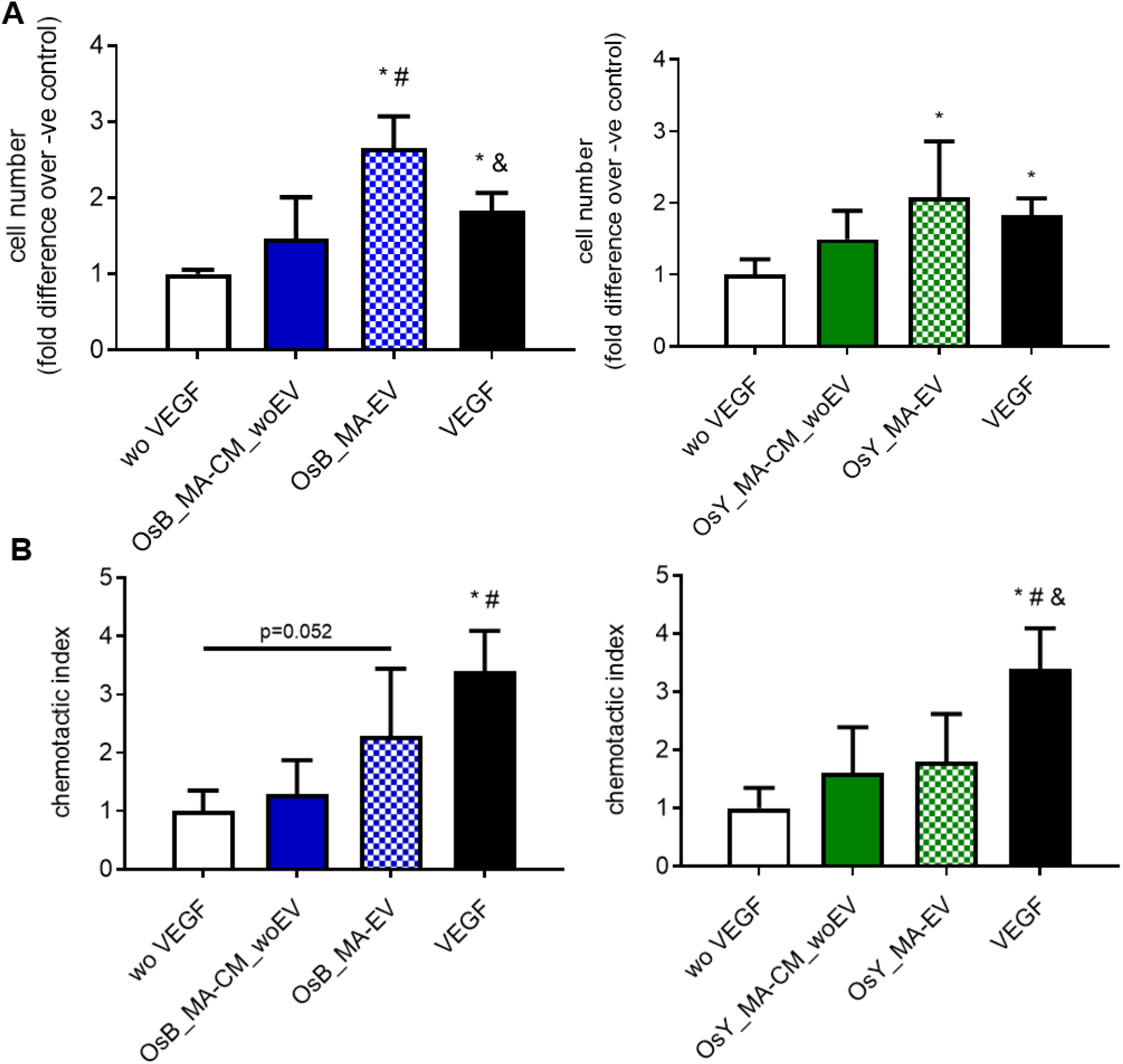
Extracellular vesicles (EVs) from mechanically activated osteoblasts and osteocytes influence HUVEC proliferation and migration. **(A)** Quantification of HUVEC number after 24 hours cultured in fresh medium without VEGF (negative control), with VEGF (positive control), mechanically activated MC3T3-E1 CM depleted of EVs (OsB_MA-CM_woEV) or isolated EVs (OsB_MA-EV), and mechanically activated MLO-Y4 CM depleted of EVs (OsY_MA-CM_woEV) or isolated EVs (OsY_MA-EV). **(B)** Quantification of HUVECs migrated through a porous membrane in fresh medium without VEGF (negative control), with VEGF (positive control), mechanically activated MC3T3-E1 CM depleted of EVs (OsB_MA-CM_woEV) or isolated EVs (OsB_MA-EV), and mechanically activated MLO-Y4 CM depleted of EVs (OsY_MA-CM_woEV) or isolated EVs (OsY_MA-EV). Data presented as Mean ± SD, N = 3-7. * p<0.05 VS wo VEGF, # p<0.05 VS MA-CM_woEV, & p<0.05 VS MA-EV.

Taken together, this data demonstrates that extracellular vesicles released from mature bone cells subjected to mechanical stimulation can enhance proliferation and, to a certain degree, the recruitment of endothelial cells in preparation for early angiogenesis.

### Mechanically stimulated osteoblasts and osteocytes secrete extracellular vesicles that promote angiogenesis

To determine whether EVs secreted by mature bone cells subjected to mechanical stimulation can coordinate angiogenesis, we next examined HUVEC tubule formation in response to EVs secreted from mechanically activated osteoblasts and osteocytes, in addition to the MA-CM depleted in EVs (Fig.6A).

**Figure 6:**
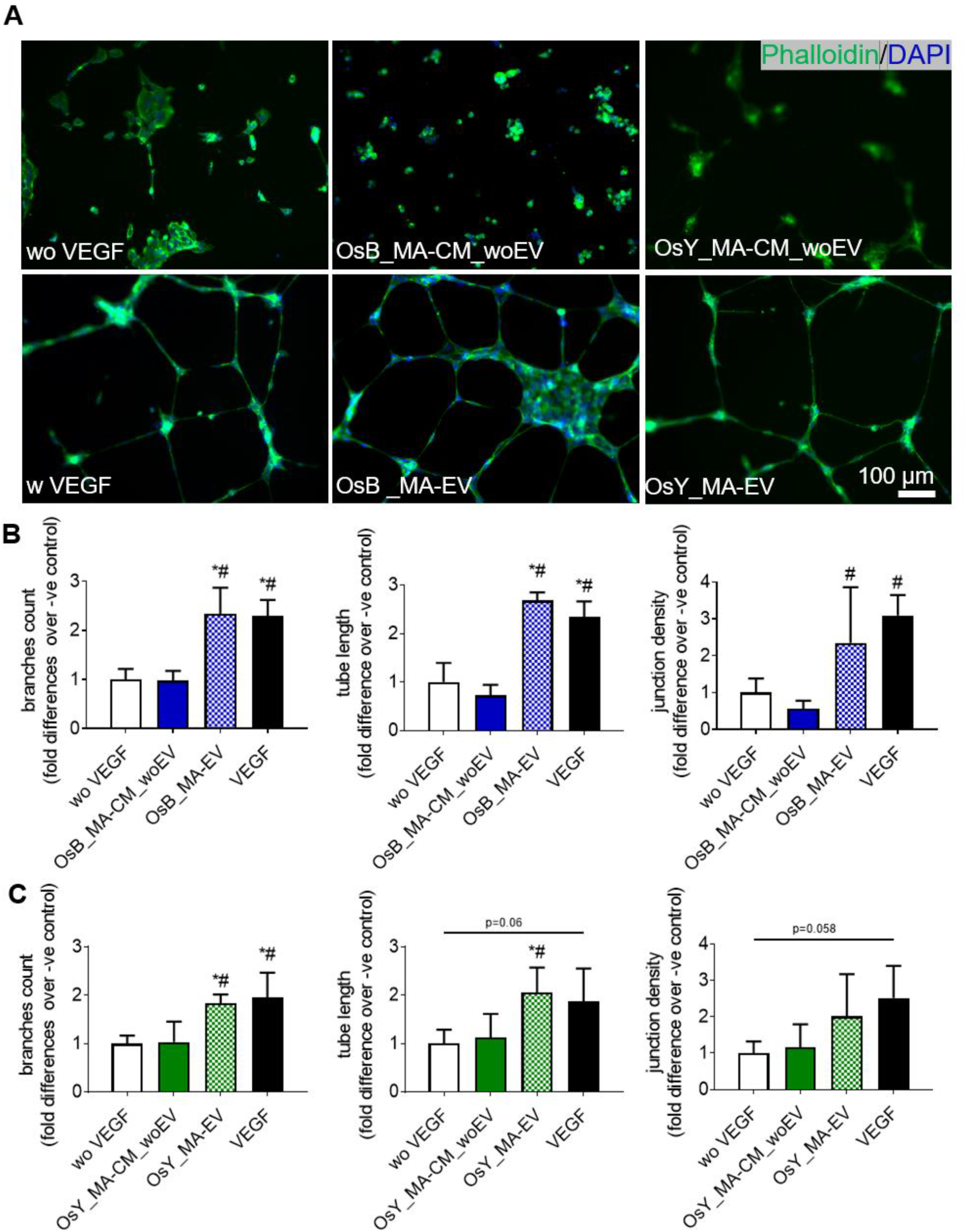
Extracellular vesicles released from mechanically activated osteoblasts and osteocytes induce vessel formation in HUVECs. **(A)** HUVECs on Matrigel in fresh medium without VEGF (negative control), with 10 ng/mL VEGF (positive control), treated with mechanically activated MC3T3-E1 CM depleted of EVs (OsB_MA-CM_woEV) or isolated EVs (OsB_MA-EV), and mechanically activated MLO-Y4 CM depleted of EVs (OsY_MA-CM_woEV) or isolated EVs (OsY_MA-EV) **(B)** Quantification of number of branches, tube length, junction density of tubes formed by HUVECs treated with groups listed above. Data presented as Mean ± SD, N = 3-7. * p<0.05 VS wo VEGF, # p<0.05 VS MA-CM_woEV, & p<0.05 VS MA-EV.

Osteoblast and osteocyte MA-CM, which previously enhanced angiogenesis, did not elicit any significant response when this media was depleted of extracellular vesicles (Fig.6A), and this was confirmed following quantification of branches, tube length, and junction density (Fig.6B,C). However, following supplementation with mechanically activated EVs isolated from osteoblast and osteocyte CM, tube formation was evident in both groups and mirrored that seen in the VEGF positive controls (Fig.6A). Furthermore, quantification of these images revealed a statistically significant increase in the number of branches, tube length, and junction density resulting from treatment with mechanical activated EVs when compared to both no VEGF negative controls and mechanically activated bone cell CM depleted of EVs (Fig.6B,C), with the exception of junction density following treatment with osteocyte derived MA-EVs. These data demonstrate that mature bone cells drive angiogenesis via an EV release mechanism following mechanical stimulation.

### Osteoblasts and osteocytes secrete low levels of VEGF, that is mechanoregulated, and independent of extracellular vesicles

VEGF is one of the most potent and widely studied proangiogenic factors that is secreted by bone cells and has been shown to be mechanically regulated [40, 41]. Therefore, to investigate whether VEGF release by osteoblasts and osteocytes may contribute to the findings of this study, we next analysed the VEGF levels in CM isolated from statically cultured and mechanically activated osteoblasts and osteocytes, in addition to MA-CM from both cells depleted of EVs and the corresponding secreted EVs.

Surprisingly, the VEGF levels in CM collected from statically cultured or mechanically activated osteoblasts was almost undetectable (<5 pg/mL; Fig.7A). While mechanical stimulation did enhance the concentration of VEGF ∼1.5-fold in the osteoblast secretome, this was not significant. Similarly, osteoblast derived MA-EVs and EV-depleted osteoblast MA-CM also contained negligible quantities of VEGF (<5 pg/mL). In contrast, osteocytes secrete higher concentrations of VEGF (67.29 ± 16.00 pg/mL; Fig.7C) and mechanical stimulation significantly enhances this 1.8-fold (126.45 ± 6.74 pg/mL; p<0.001). Interestingly, almost no VEGF was detected in osteocyte derived MA-EV with all VEGF being detected in MA-CM depleted of EVs (99.26 ± 17.25 pg/mL), indicating that VEGF is not specifically packaged in EVs prior to cellular release.

**Figure 7:**
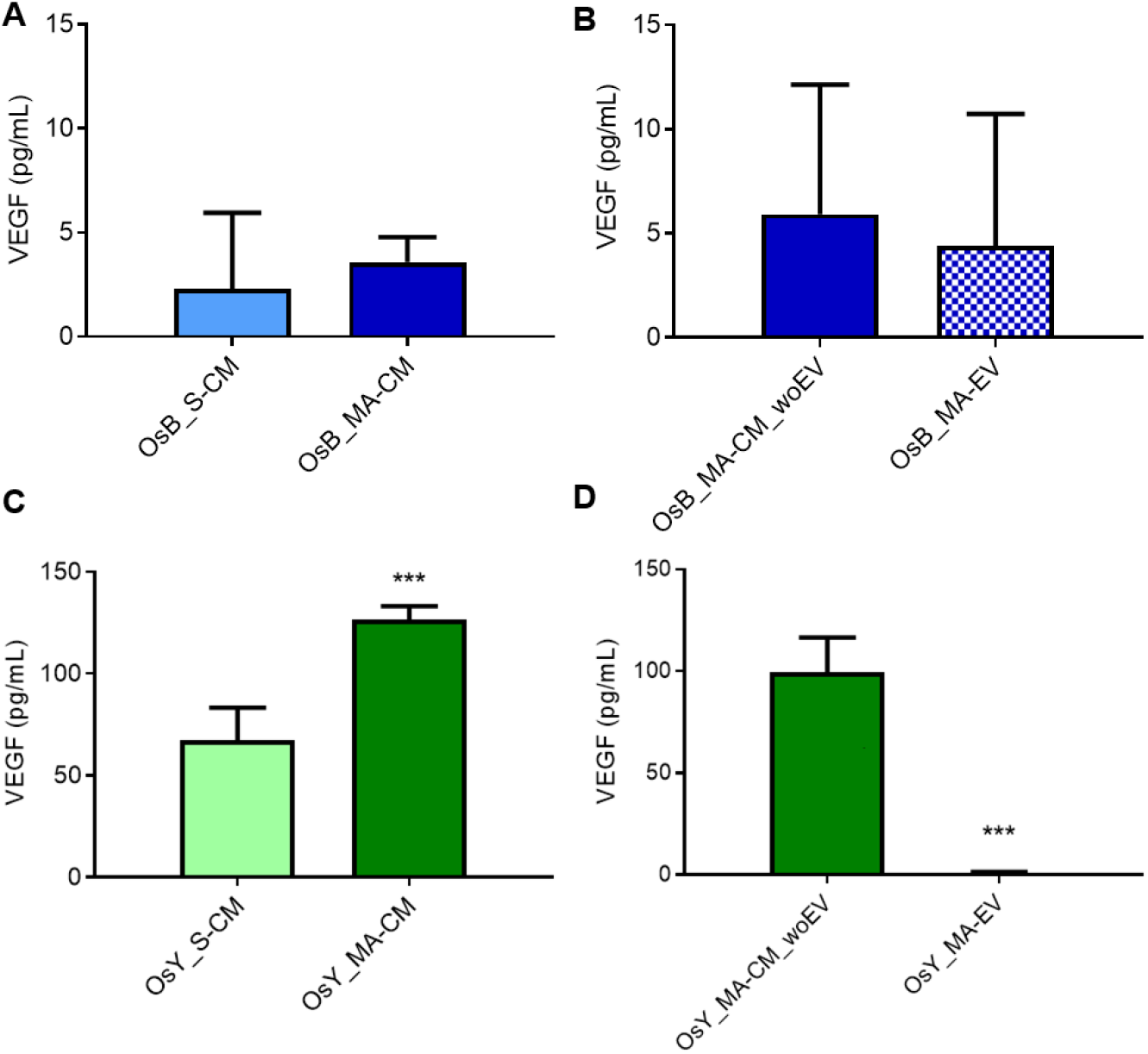
Osteoblasts and osteocytes secrete low levels of VEGF that is mechanoregulated and independent of extracellular vesicles. VEGF protein expression within **(A)** statically and mechanically activated osteoblast conditioned media (CM), **(B)** mechanically activated osteoblast CM depleted of EVs and osteoblast derived MA-EVs. VEGF protein expression within **(C)** statically and mechanically activated osteocyte conditioned media (CM), **(D)** mechanically activated osteocyte CM depleted of EVs and osteocyte derived MA-EVs. Data bars indicate mean ± SD, N = 6. *** p<0.001.

Taken together, this data demonstrates that the mature bone cells, particularly osteocytes, secrete the pro-angiogenic factor VEGF and that this secretion is mechanically regulated but not packaged within extracellular vesicles. Compared to the positive control utilized in this study to successfully induce angiogenesis (10 ng/mL), the VEGF levels detected in all groups was still relatively low and unlikely to be responsible for angiogenic effects identified.

### Osteocyte release extracellular vesicles that contain miRNAs associated with angiogenesis that are increased with mechanical stimulation

As extracellular vesicles have been shown to mediate communication via the packaging and delivery of miRNAs [42], many of which are known regulators of angiogenesis [43], we next looked to profile the miRNA cargo within MA-EVs.

miRNAs within extracellular vesicles released from statically cultured and mechanically stimulated osteocytes were profiled in three replicates using high throughput sequencing (miRNA-seq). A total of 550 known (mature) murine miRNAs were detected in EVs collected from both groups. Of these, 428 (78%) were classified as lowly expressed and removed from the analysis. The remaining 122 miRNAs showed a strong expression signal and were used in downstream analysis. Functional overrepresentation analysis (ORA) was performed on this set of miRNAs using miEAA, a web-based application. Over-representation analysis found a significant enrichment of miRNAs associated with gene ontology categories such as blood vessel remodelling, cell proliferation, and positive regulation of angiogenesis (Fig.8A). Differential expression analysis using DESeq2 revealed two distinct miRNAs that were significantly upregulated in MA-EVs when compared to EVs derived from statically culture osteocyte conditioned media (mmu-miR-150-5p (log2 fold change = 2.3; p-adjusted value = 0.01) and mmu-miR-2137 – log2 fold change = 2.3; p-adjusted value = 0.02) (Fig.8B,C). A total of 556 target genes were mapped to these two differentially expressed miRNAs through TargetScan analysis. Functional enrichment analysis identified 13 significant KEGG pathways and 380 significant GO terms derived from these target genes. Fig. 8D,E show a selection of enriched categories which are significantly associated with angiogenesis and related terms.

**Figure 8:**
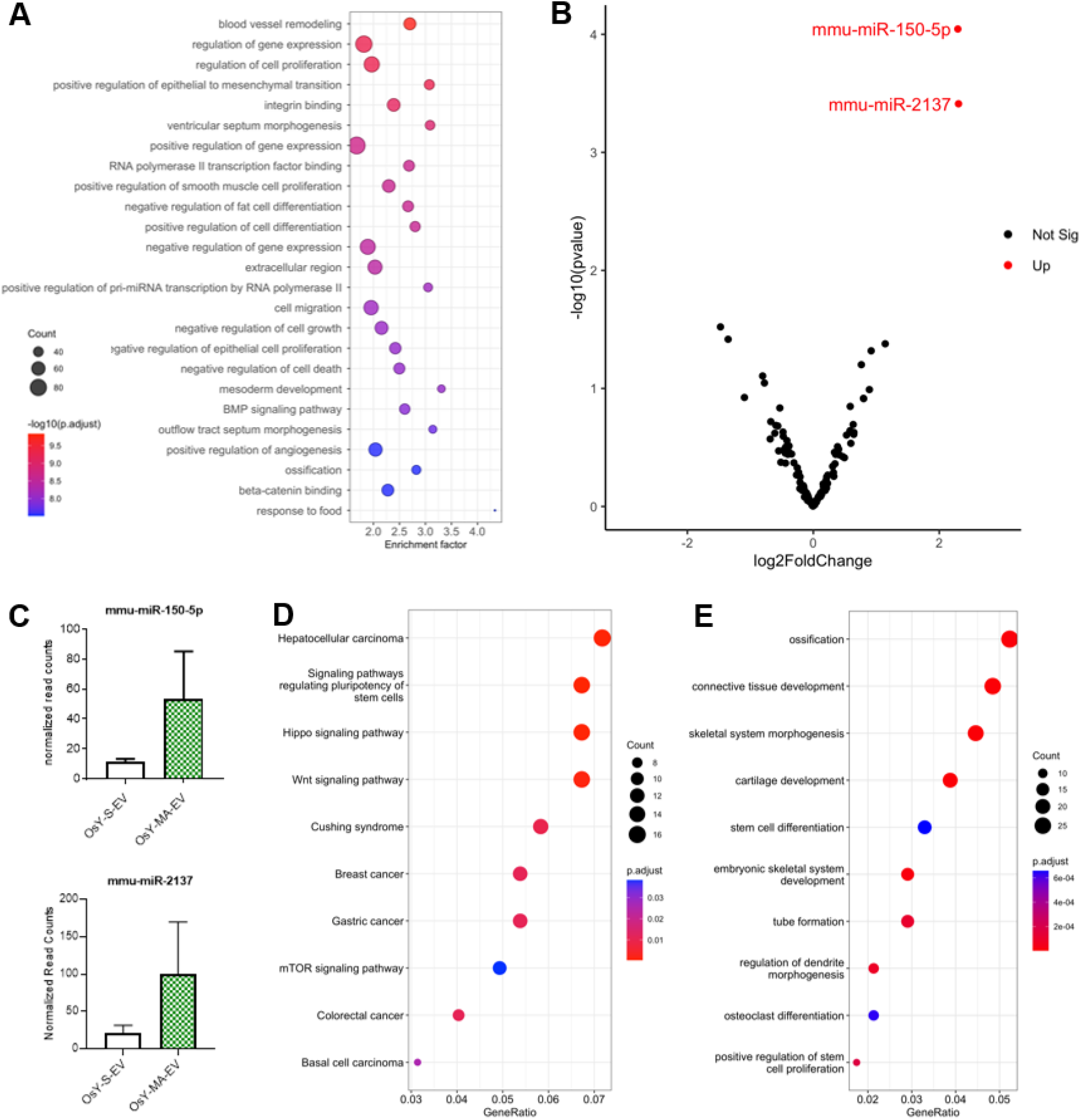
Osteocyte release extracellular vesicles that contain miRNAs associated with angiogenesis that are increased with mechanical stimulation. **(A**) miEAA Enrichment analysis of the 122 miRNAs detected in static (S-EV) and mechanically activated osteocyte derived EVs (MA-EV). Top 25 GO terms which include some associated with blood vessel remodelling, cell proliferation and angiogenesis. **(B)** Volcano plot of miRNAs expressed in S-EVs vs MA-EVs. Red dots represent the upregulated miRNAs (fold change >= 2; p-adjusted value <= 0.05). **(C)** Expression barplots of miR-150-5p and miR-2137. **(D-E)** KEGG and GO enrichment analysis of the targeted genes derived from TargetScan predictions of both miR-150-5p & -2137.

## Discussion

Blood vessel formation is an important initial step for bone formation during development as well as during remodelling and repair in the adult skeleton. This results in a heavily vascularized tissue where endothelial cells and skeletal cells are constantly in crosstalk to facilitate homeostasis, a process that is mediated by numerous environment signals, including mechanical loading. Breakdown in this communication can lead to disease and/or poor fracture repair. Therefore, this study aimed to determine the role of mature bone cells in regulating angiogenesis, how this is influenced by a dynamic mechanical environment, and understand the mechanism by which this could occur. Herein, we demonstrate that both osteoblasts and osteocytes can coordinate endothelial cell proliferation, migration, and vessel formation via a mechanically dependent paracrine mechanism. Moreover, we identified that this process is mediated via the secretion of extracellular vesicles as isolated EVs from mechanically stimulated bone cells elicited the same response as that seen with the full secretome. Lastly, despite mechanically activated bone cell derived EVs driving a similar response to VEGF treatment, MA-EVs contain minimal quantities of this angiogenic factor, indicating that this EV mediated angiogenic response is not dependent on the secreted VEGF protein. Lastly, we profiled the miRNA cargo of MA-EVs and identified two miRNAs that are mechanically regulated and associated with angiogenesis, indicating that the pro-angiogenic effect of bone cell derived MA-EVs may be mediated by an RNA-based mechanism. Taken together, this study highlights an important mechanism in the osteogenic-angiogenic coupling in bone that is present only with mechanical loading and has identified the mechanically activated bone cell derived extracellular vesicle as a potential therapeutic to promote angiogenesis.

Osteoblasts and osteocytes can coordinate angiogenesis via a mechanically driven paracrine mechanism. Previous work by Liu *et al*. has shown that fluid shear can regulate angiogenesis in bone by modulating osteoblast and endothelial crosstalk [2]. We confirmed that shear stress stimulation of osteoblasts does indeed potentiate angiogenesis, in addition to endothelial cell proliferation and migration. While osteocytes are the most abundant cell type in bone and are known to play a role in transducing mechanical signals in bone, their role in regulating bone angiogenesis has seldom been studied. Prasadam *et al*. showed that the osteocyte lacuna-network was intimately associated with the blood vessels and the osteocyte dendrites were directly connected with the vessel wall in bone matrix [15]. This suggests a possibility of a close interaction of osteocytes with endothelial cells in bone. In this study, we have shown that osteocytes release paracrine signals that can regulate angiogenesis in response to shear stress. The fluid shear stress was applied via an orbital shaker system. The average shear stress produced by this system is 0.32 Pa, and the maximal shear stress is approximately 1.3 Pa [33]. This range is comparable to the lower stimulatory range for osteoblasts and osteocytes [44]. Estimation of shear stress values that osteoblasts are exposed to is complicated, however, based on experimental and computational studies, it was hypothesized that the stress regime osteoblasts experience is distinctively different to osteocytes and it might encounter lower, interstitial-like shear stress [45, 46]. This demonstrates that the low fluid shear stimulus is sufficient to drive the release of pro-angiogenic factors from mature bone cells, highlighting a potential mechanism of osteogenic-angiogenic coupling in bone mechanobiology.

Osteoblasts and osteocytes can coordinate angiogenesis in response to mechanical stimulation via the release of extracellular vesicles. Previous studies have demonstrated that EVs released from osteoblasts and osteocytes can induce MSC osteogenic differentiation [23, 47, 48], and that this is enhanced following mechanical stimulation of the osteocyte [30, 31, 49]. Moreover, osteoblast-derived exosomes contain osteoprotegerin, RANKL and tartrate-resistant acid phosphatase, which are critical for osteoclast differentiation [50]. Other studies have indicated that mesenchymal stem cell-derived EVs contain a set of angiogenic factors such as interleukin-8 and miRNAs, which can significantly promote endothelial cell proliferation and tube formation [51, 52]. Despite these promising results, few studies have focused on the role of osteoblast EV or osteocyte EV on angiogenesis. We demonstrated that mature bone cell derived EVs can enhance endothelial cell proliferation, migration and vessel formation to a similar extent to that seen with cells treated with VEGF. This therefore highlights the mechanically stimulated bone cell-derived EV as a potential pro-angiogenic therapeutic.

Although increased VEGF was detected in mechanical activated osteoblast and osteocyte CM when compared to static CM, the level of the VEGF is relatively low (pg range), and thus is not likely to be contributing to the angiogenic response to bone derived MA-EVs. Consistent with the results of our study, Thi *et al* detected that osteoblasts release VEGF at a level approximately 18 pg/μg of protein in response to parallel flow shear stress [40] and Liu *et al*. did not detect VEGF in osteoblast CM [2], postulating that this may be due to the released VEGF attaching to the extracellular matrix [2]. Interestingly, we detected a lower level of VEGF in bone derived MA-EVs compared to the full CM. Similarly, MSC-derived EVs have proven angiogenic properties, however, MSC-derived EVs contain significantly lower levels of VEGF than full CM [52]. VEGF is known to stimulate angiogenesis in a strict dose-dependent manner and therefore the low concentration of VEGF present in the CM or EVs might be insufficient to induce angiogenesis [53]. Therefore, VEGF release may not be the mechanism underlying this EV based osteogenic-angiogenic coupling. This might suggest a possible role for other angiogenic factors such as PDGFAA, OPN, or CXCL12 which are known to be released by mature bone cells in response to mechanical stimulation [2, 54]. Moreover, non-protein-based cargo such as RNAs, specifically miRNAs, may mediate this angiogenic effect of MA-EVs [52, 55]. Both miRNA-150-5p and miRNA-2137 expression were found to be significantly increased within MA-EVs when compared EVs collected form statically cultured osteocytes. While little is known about the angiogenic properties of miR-2137, miR150-5p is strongly associated with angiogenesis. For example, miR-150 secreted by monocytes has been shown to induce endothelial tube formation *in vitro* and angiogenesis *in vivo*, and down-regulation of miR-150 has been linked to an inhibition of angiogenesis seen in diabetes, cancer, and atherosclerosis [56]. Fang *et al*. elucidated the exosomal miRNA expression profile during osteonecrosis of femoral head (ONFH) and identified that miR-150-5p was significantly downregulated in exosomes within the plasma of the femoral head. Moreover, they demonstrated that miR-150-5p-modified MSC exosomes can promote angiogenesis potentially protecting against ONFH [57]. The target genes of miR-150-5p participate in the TGFβ, MAPK, HIF-1, PI3K-Akt, and mTOR signalling pathways, all of which been linked to angiogenesis. Therefore, miRNAs such as miR-150-5p are also likely to play a role in this mature bone cell EV regulation of angiogenesis. Future work will focus on identifying and validating the angiogenic cargo within bone derived MA-EVs.

In conclusion, this study demonstrates a novel mechanism by which mechanical loading regulates blood vessel formation via the release of extracellular vesicles from mature bone cells such as the osteoblast and osteocyte. This opens up a new avenue to study the potential of mechanically activated bone cell derived extracellular vesicle as a novel therapeutic to promote angiogenesis, with applications in bone regeneration.

## Author Contributions Statement

N.S., M.M., and D.H. Study conceptualisation, data analysis, prepared figures, and wrote the main manuscript text. N.S., K.E., E.S., I.W., M.L., F.M.R., K.H., L.O’D. Data collection and analysis. All authors contributed with critical revisions and editing of the manuscript.

## Data availability

The datasets generated during and/or analysed during the current study are available from the corresponding author on reasonable request.

## Acknowledgement

The authors would like to acknowledge funding from Science Foundation Ireland (SFI) Frontiers for the Future Grant SFI 19/FFP/6533 and the Irish Research Council Advanced Laureate Award EVIC [IRCLA/2019/49], and Horizon 2020 Research and Innovation Award EVPRO [814495]. We thank Dr Svenja Sladek and Prof. Lidia Tajber for their help with Nanoparticle tracking analysis. We also appreciate the help from Mr. Neal Leddy with transmission electron microscopy.

